# Large-Scale Computational Modeling of H5 Influenza Variants Against HA1-Neutralizing Antibodies

**DOI:** 10.1101/2024.07.14.603367

**Authors:** Colby T. Ford, Shirish Yasa, Khaled Obeid, Rafael Jaimes, Phillip J. Tomezsko, Sayal Guirales-Medrano, Richard Allen White, Daniel Janies

## Abstract

The United States Department of Agriculture has recently released reports that show samples from 2022-2024 of highly pathogenic avian influenza (H5N1) have been detected in mammals and birds (1). To date, the United States Centers for Disease Control reports that there have been 27 humans infected with H5N1 in 2024 (2). The broader potential impact on human health remains unclear. In this study, we computationally model 1,804 protein complexes consisting of various H5 isolates from 1959 to 2024 against 11 hemagglutinin domain 1 (HA1)-neutralizing antibodies. This study shows a trend of weakening binding affinity of existing antibodies against H5 isolates over time, indicating that the H5N1 virus is evolving immune escape of our medical defenses. We also found that based on the wide variety of host species and geographic locations in which H5N1 was observed to have been transmitted from birds to mammals, there is not a single central reservoir host species or location associated with H5N1’s spread. These results indicate that the virus has potential to move from epidemic to pandemic status in the near future. This study illustrates the value of high-performance computing to rapidly model protein-protein interactions and viral genomic sequence data at-scale for functional insights into medical preparedness.

**Research in Context:** *Evidence before this study:* Previous studies have shown cases of avian influenza transmissions to mammals that are increasing in frequency, which is of concern to human health. Since 1997, nearly a thousand H5N1 cases have been reported in humans with a 52% fatality rate. Previous analyses have indicated specific mutations on the hemagglutinin protein that allow for this “host jumping” between birds and mammals (3). There are also existing evidence of recent viral strains with reduced neutralization to sera (4).

*Added value of this study:* This study provides a comprehensive look at the mutational space of hemagglutinin of H5N1 influenza and presents computational predictions of the binding between various HA1-neutralizing antibodies derived from infected vaccinated patients and humanized mice and 1,804 representative H5 HA1 proteins. These analyses show a weakening trend of existing antibodies. We also confirm that the mutations found in other studies that enable zoonosis also affect binding affinities of the antibodies tested. Furthermore, through phylogenetic analyses, we quantify the avian-to-mammalian transmissions from 1959 to 2024 and show a persistent circulation of isolates between North America and Europe. Taken together, the continuous transmission of H5N1 from birds to mammals and the increase in immuno-evasive HA strains in mammals sampled over time suggest that antigenic drift is a source of spillover risk.

*Implications of all the available evidence:* Our findings indicate that the worsening in antibody binding, along with the increase in of avian-to-mammalian H5N1 influenza transmissions are risks to public health. Through the findings of previous studies along with the predictions reported in this study, we can now monitor specific mutations of interest, quantified by their potential impact on antibody evasion, and inform public health monitoring of circulating isolates in 2024 and beyond. In addition, these findings may help to guide future vaccine and therapeutic development in the fight against H5N1 influenza infections in humans.

## Introduction

Wild aquatic birds (e.g., order Anseriformes) are fundamental hosts of influenza viruses, which are transmitted from domestic birds (e.g., Galliformes) and mammals (orders Artiodactyla, Carnivora, and Primates) (5, 6). H5N1 has circulated in nature since 1959, following an outbreak in Scotland in chickens (7). In 1996, H5N1 influenza largely occurred in Anseriformes and spread to humans and chickens in Hong Kong in 1997 (6, 8). As a response to H5N1 in chickens and occasional human infection, chickens were culled in Hong Kong in the period 1997 to 2011 (9). In 2002, the most common hosts of H5N1 were Anseriformes with occasional transmission to Galliformes and humans throughout China and South East Asia (6, 10). Since 2003, various lineages of H5N1 have spread throughout China and Hong Kong, South East Asia, Russia, North Africa, the West Bank, Gaza Strip and Israel, Pakistan, Bangladesh, India, Bhutan, Nepal, Europe, Japan and South Korea, using a wide variety of hosts (11, 12). Many avian taxa (Charadriiformes, Accipitriformes, Corvidae, Ardeidae, Columbidae, and Passeriformes) as well as primate, carnivore, artiodactyl, and arthropod hosts have been infected with H5N1 (6). H5N1 infections in humans have been reported by the World Health Organization (WHO) in: Hong Kong 1997–2003, in China and Hong Kong 2003–2014; in Thailand 2003–2007; in Indonesia 2005–2012; in Nigeria in 2007; in Bangladesh 2011-2013; in Azerbaijan, Turkey, Iraq, Myanmar, Pakistan, and Djibouti in 2006-2007; in Egypt 2003-2014; in Lao PDR 2007; in Vietnam 2003–2014; in Cambodia 2003–2014, and in Canada in 2014. The document produced by the WHO has not been updated since (13).

However, a comprehensive review from 2023 (14) illustrates the recent (2022-23) spread (on top of the previous spread) of H5N1 in animals as follows:

- **Asia:** Bhutan, Hong Kong, India, Japan, Korea, Nepal, Philippines, Taiwan, and Vietnam.
- **Europe:** Albania, Austria, Belgium, Bosnia and Herzegovina, Bulgaria, Croatia, Czechia, Denmark, Estonia, Finland, France, Germany, Greece, Hungary, Iceland, Ireland, Italy, Latvia, Lithuania, Luxembourg, Moldova, Montenegro, Netherlands, North Macedonia, Norway, Poland, Portugal, Romania, Russia, Serbia, Slovakia, Slovenia, Spain, Sweden, Switzerland, and the United Kingdom.
- **The Middle East:** Israel and Turkey.
- **Africa:** Algeria, Gambia, Nigeria, Reunion, Senegal, and South Africa.
- **North and South America:** Bolivia, Brazil, Canada, Bolivia, Chile, Colombia, Ecuador, Guatemala, Honduras, Mexico, Panama, Peru, Venezuela, and the United States.

In 2024, H5N1 has been found in animals in Antarctica (15). Also, in March 2024, an outbreak of H5N1 was reported among several herds of U.S. dairy cattle. H5N1 also caused fatal infections among cats, infection in poultry, and four reported infections in dairy workers (16–18). From 1997 to late April 2024, 909 human H5N1 cases were reported, with 52% of cases being fatal (19).

Continued transmission of avian strains of H5N1 to livestock and humans may lead to subsequent human-to-human transmission which can devastate public health world-wide. As the human-animal interface increases due to shrinking natural habitats, deforestation, and increased demand for animal products; animal-human disease transmission is becoming more common (20).

Current human seasonal influenza vaccines do not confer immunity against H5N1 influenza or other animal influenza A viruses (21). Moreover, recent studies have shown there is little existing immunity to H5N1 in the USA (22). Such immunity may exist elsewhere in the world and in populations in due to previous infection or immunization.

Thus, it is of great public health interest to discover rapidly develop molecular insight into the effects of mutations of H5N1 on existing human immunity (23). In this study, we present the results of a large computational corpus of molecular docking experiments between various H5 isolates against existing HA1-neutralizing antibodies and show changes over time.

## Methods

This study closely follows the published methods in Tomezsko and Ford et al. (2024) and other protein modeling studies by this team (25–29). The specific workflow for this study is depicted in Figure 1.

**Fig. 1.**
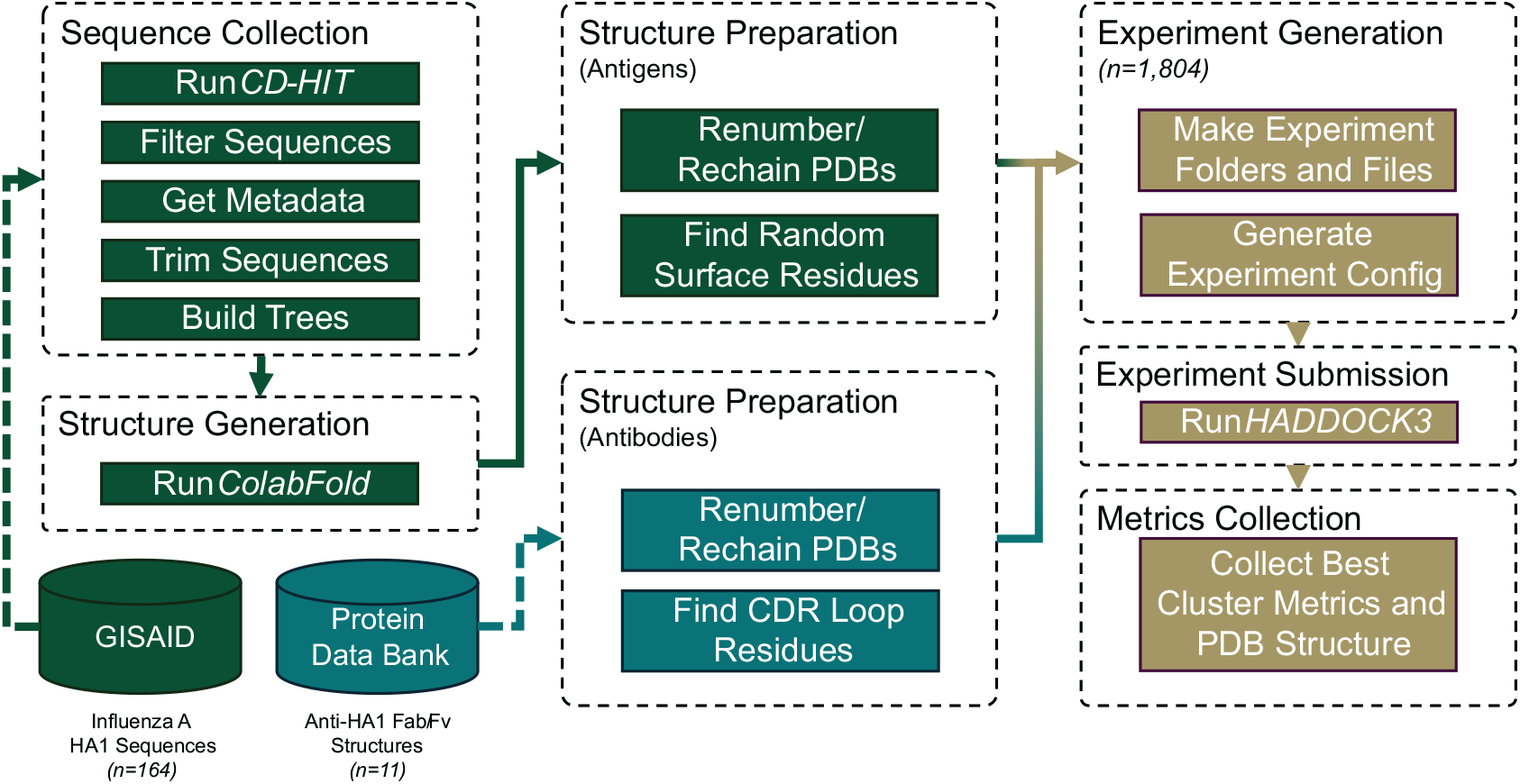
Workflow diagram of the data procurement, data preparation, and analysis steps.

### HA1 Sequence Collection

18,693 influenza A H5 sequences were downloaded from the GISAID EpiFlu database (30, 31) on June 17, 2024. Isolate metadata, including: date of viral isolation, country of origin, and host information, were also collected. We then derived taxonomic classifications from the provided host metadata along with continent of origin from the country information.

These amino acid sequences were clustered using CD-HIT v4.8.1 (32, 33). Ranging in size from 11 to 576 amino acids, the process resulted in 250 clusters, based on 97% identity. A representative sequence from each of these clusters was selected, which was then output in a FASTA file.

These representative sequences were then aligned to each other with MUSCLE v3.8.425 (34) using default settings. Then, sequences that were of low quality, incomplete, or did not contain the desired hemagglutinin domain 1 (HA1) region were removed. This resulted in 164 amino acid sequences that were used in the subsequent steps. These sequences were trimmed to the HA1 receptor binding domain (approximately residues 111-269, depending on the presence of indels) (35).

Also, clusters consisting of only lab derived isolates (*n* = 3) were analyzed, but have been left out of the reported statistics and visualizations.

### Structure Prediction

Structures for each of the 164 HA1 sequences were predicted using ColabFold v1.5.5 (36), a protein folding framework that uses AlphaFold2 (37) accelerated with MMseqs2 (38), with default settings. The side chains of these predicted structures were relaxed using the OpenMM/Amber method (39). The .PDB file output of the top structure (i.e., the one with the highest pLDDT^1^ confidence) was selected and used for subsequent analyses.

### Neutralizing Antibody Collection

Existing structures for 11 HA1-neutralizing antibodies were collected by a database search through Thera-SAbDab (40) and the Protein Data Bank (41). Each of these 11 antibody structures have an epitope on the target HA1 domain of the hemagglutinin protein, though not all share the same epitope. These antibodies and their respective PDB IDs are listed in Table 1.

**Table 1.**
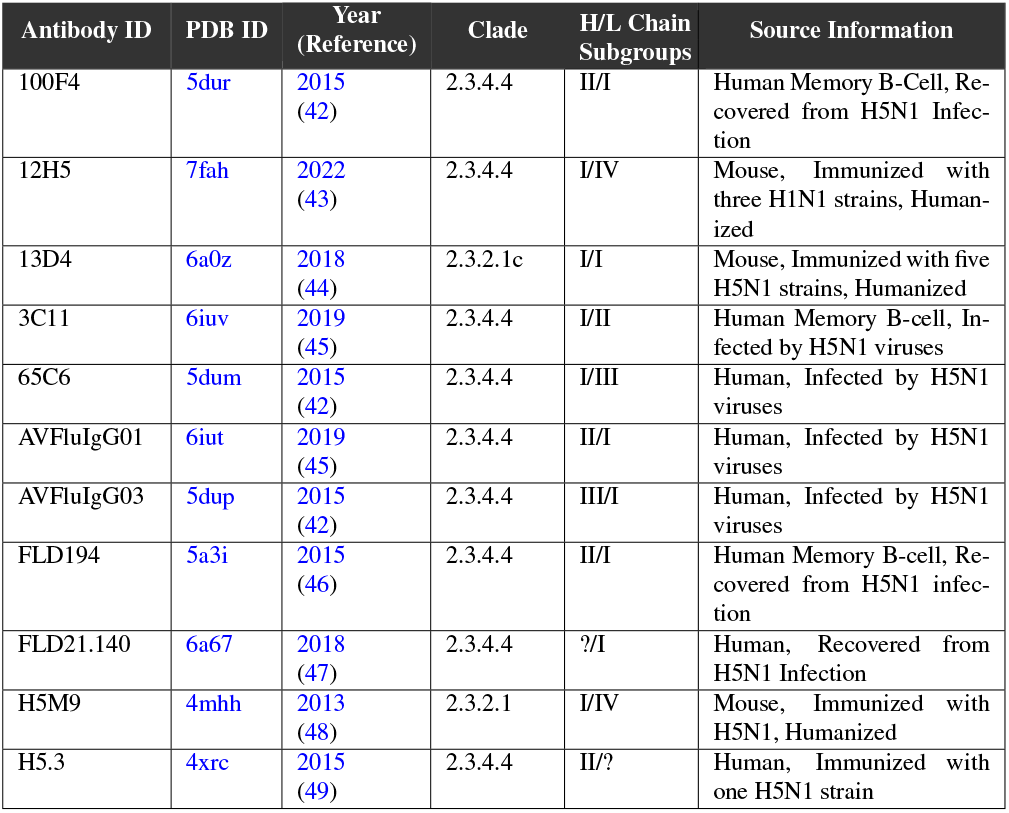
HA1-neutralizing antibodies.

### Docking Analyses

Using HADDOCK3, a computational framework for the integrative modeling of biomolecular complexes (50), each antibody was docked to each antigen across the dataset. Given the 11 antibody structures and 164 HA1 structures, this resulted in 1,804 docking experiments to be performed.

#### Experiment Generation

HADDOCK3 inputs for each experiment were generated programmatically, defining the antibody and antigen .PDB file inputs on which to dock. Other experiment files were also copied or created programmatically including the scripts to run the docking process and to generate other configuration files.

HADDOCK3 requires the definition of active and inactive residue restraints (AIRs) to help guide the protein docking process. To avoid biasing the docking placement of the antibody on the HA1 antigens, a random subset of surface residues were selected as “active” and were then included in the AIR file on which to dock. For the antibody structures, residues in the CDR loops were detected using ANARCI (51), a Python package for numbering antibody sequences.

Lastly, HADDOCK3 configuration files were generated programmatically, which define the input .PDB files, the output directory, and the steps of the docking process. The logic for this programmatic generation of HADDOCK3 configuration files is available in the supplementary GitHub repository.

### Docking Process

HADDOCK3 provides a configurable interface for defining the individual steps of the docking process, including the rigid-body docking, flexible refinement, and solvent-based refinement, along with any desired clustering and filtering steps.

For this study, we customized the published HAD-DOCK3 protocol for antibody-antigen modeling (52) to focus on generating the singular best cluster of docking results for each experiment and reducing excess work by the docking process. The specific steps of our HADDOCK3 configuration that was used for the experiments are shown in Table 2.

**Table 2.**
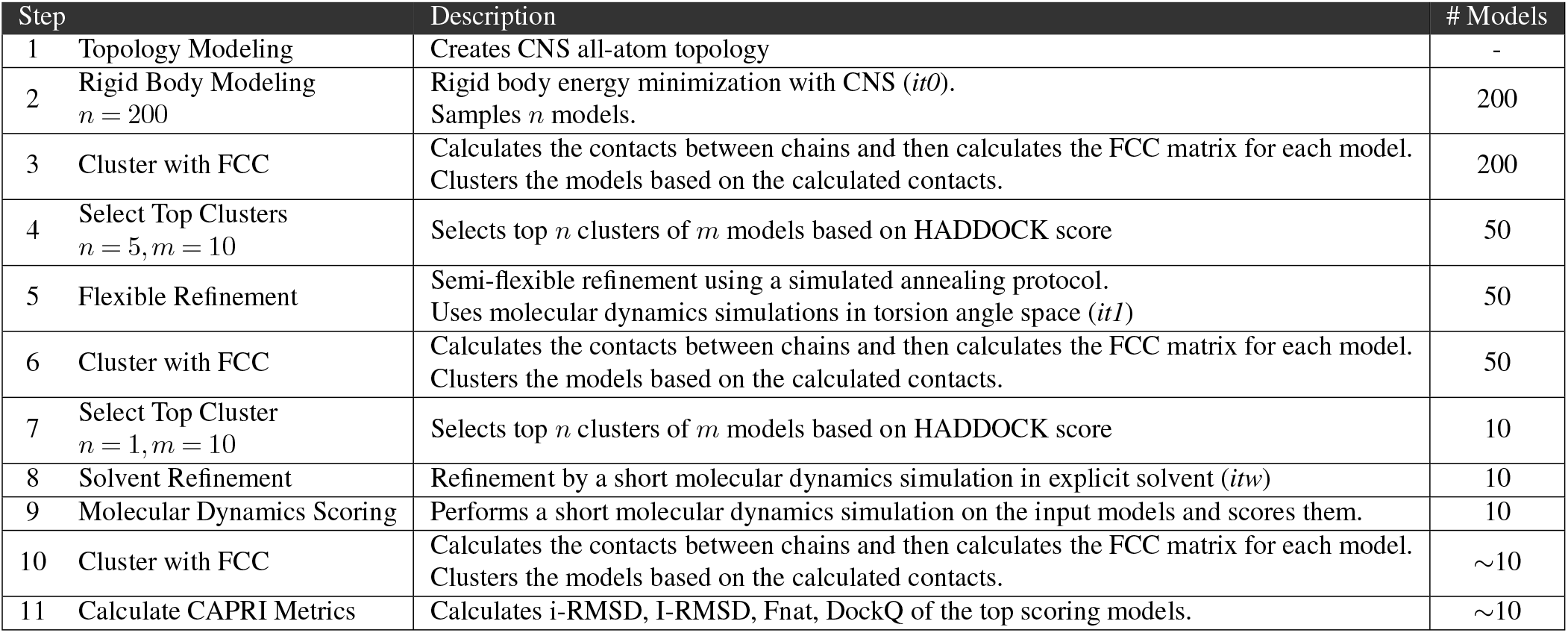
Descriptions of the steps of the HADDOCK3 docking configuration.

The HADDOCK3 system outputs multiple metrics for the predicted binding affinities and an output set of PDB files containing the antibody docked against the HA1 antigen. Some main metrics include:

- Van der Waals intermolecular energy (*vdw*)
- Electrostatic intermolecular energy (*elec*)
- Desolvation energy (*desolv*)
- Restraints violation energy (*air*)
- Buried surface area (*bsa*)
- Total energy (*total*): 1.0*vdw* + 1.0*elec*
- HADDOCK score: 1.0*vdw* + 0.2*elec* + 1.0*desolv* + 0.1*air*

Note that the HADDOCK Score is a conglomerate metric used to assess the best complexes (or best cluster of complexes) that get promoted through the various refinement iterations in the pipeline.

#### Computational Scalability

For this study, we used a Docker containerized version of HADDOCK3^2,3^, which contains all of the software dependencies to allow HADDOCK3 to run more readily in a high-performance computing (HPC) environment.

HADDOCK3 was run in a Singularity container on the UNC Charlotte Orion HPC cluster on 14 nodes, each with dual 18-core Intel Xeon Gold 6154 3.00GHz CPUs (36 cores per node). The average walltime of the experiments was 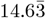 minutes. The entire set of 1,804 experiments was completed in under 2 days.

#### Post Processing

Once all experiments were completed, the metrics for each experiment were either retrieved from the CAPRI evaluation outputs (if the FCC clustering algorithm (53) reached convergence) or from the REMARK entries of the best cluster’s .PDB files. These metrics were organized in a single aggregate table, representing each experiment’s best cluster metrics, for subsequent visualization and statistical analyses. The full table of experiment results is available in the Supplementary Data.

### Phylogenetic, Statistical, and Protein Structure Analyses

For phylogenetic analyses, laboratory derived isolates were filtered out (*n* = 178, shown in Table 3), resulting in a set of 18,515 isolates. An alignment 18,515 HA sequences was generated using MAFFT v7.471 (54) under default settings. Next, a phylogenetic tree search was performed using this alignment with TNT v1.6 (55) using the commands: xmult= level 1 rep 1000. One of the best scoring trees was used for downstream analyses. StrainHub v2.0.0 (56) was used to generate transmission networks of the phylogenetic tree by host class and continent in Figure 2.

**Table 3.** Taxonomic breakdown of the isolates used in this study.

| Class | Order | Isolates | % of Total |
| --- | --- | --- | --- |
| Aves | Galliformes | 5,844 | 31.26% |
|  | Anseriformes | 4,469 | 23.91% |
|  | Other Orders | 934 | 5.00% |
|  | Not Specified | 5,287 | 28.28% |
|  | *Aves Total* | *16,534* | *88.45%* |
| Mammalia | Primates (Humans) | 666 | 3.56% |
|  | Carnivora | 191 | 1.02% |
|  | Artiodactyla | 57 | 0.30% |
|  | Other Orders | 6 | 0.03% |
|  | Not Specified | 360 | 1.93% |
|  | *Mammalia Total* | *1,280* | *6.85%* |
| Insecta | *Insecta Total* | *3* | *0.02%* |
| Other | Laboratory Derived | 178 | 0.95% |
|  | Other / Environmental | 698 | 3.73% |
|  | *Other Total* | *876* | *4.69%* |
| **Grand Total** |  | **18,693** | **100%** |

**Fig. 2.**
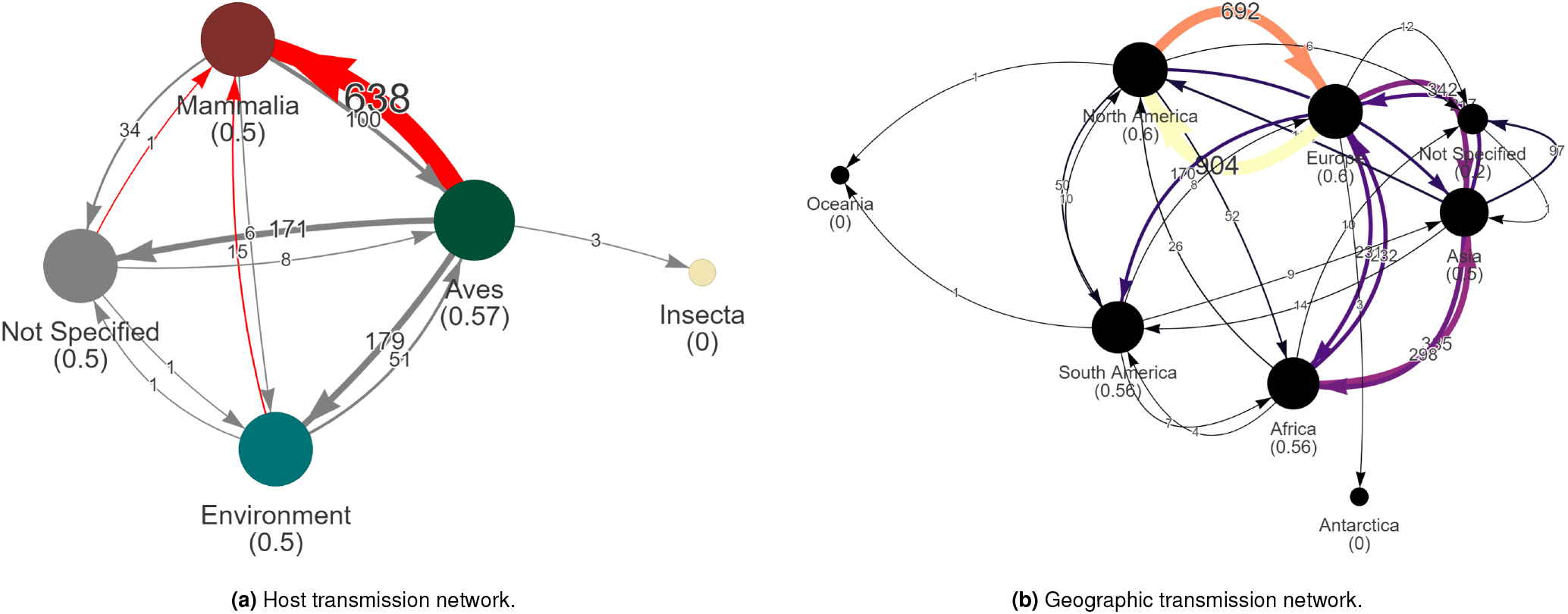
Transmission networks, generated by StrainHub, showing transmissions between (a) hosts and (b) continents. Node values in parentheses represent the source-hub ratio of that class or location. A source-hub ratio of 1 indicates that the state is always the source of the transmission. The numerical labels annotated on the edges of the graphs represent the number of transmissions as seen across the phylogenetic tree’s branches as measured in changes in metadata states.

Statistical analyses were performed using R v4.3.4 (57) and plots were generated using ggplot2 (58) and ggpubr (59). Any statistical significance reported in this study is based on a p-value threshold of *α <* 0.05.

Visualizations and analyses of the protein complexes were generated using PyMOL v2.4.1 (60) and BioPandas v0.4.1 (61).

#### Graph-based Interface Residue Assessmen

Graph-based interface residue assessment function (GIRAF) was employed to evaluate the evolution of the binding pocket with each antibody as previously described (24). The outgroup was first selected as sequence EPI242227, and a graph was computed based on the interface residues with each antibody to generate reference complexes. Subsequent graphs were then computed for each strain sequence and antibody pair. The graph edit distance (GED) was computed as the number of edits to the strain:antibody complex from the reference outgroup:antibody complex. Substitutions, deletions, and additions, were all equally weighted as a value of “1”.

### Ethics Statement

No human or animal samples were used in this study. This study was conducted in accordance with the data usage guidelines of GISAID and the research ethics policies of the University of North Carolina at Charlotte.

## Results

From the study set of 18,693 H5 isolates, we show a break-down of hosts similar to what has been reported in previous studies, indicating the representative nature of our curated dataset (5, 6). As shown in Table 3, approximately 88% of the isolates are from birds (class Aves). Also, note that all 666 isolates from Primates were collected from humans.

As shown in Supplementary Figure 2, the proportion of H5 isolates collected from various continents has changed over time. To date, isolates collected in 2024 are predominately from Europe.

### Sequence Analyses

The clustering of the 18,693 HA1 sequences resulted in 250 distinct clusters at *≥*97% identity. Further organization of the representative sequences of each cluster, shown in Supplementary Figure 3, indicates a continuous distribution of antibody binding performance.

Of note, antibodies 12H5, 3C11, 65C6, AVFlulgG01, and H5M9 show the best performance overall, though there are exceptions of poor performance (e.g., 12H5’s interaction with EPI893474 has a poor Van der Waals energy of -39.93).

#### Phylogenetic and Transmission Network Analyses

Specifically, the most common pattern is avian-to-mammalian transmission, of which there are over 600 events across the tree (see Figure 2a). There are also frequent transmission events between many continents. Bidirectional transmissions between Europe and North America are very frequent as shown by the thick orange and yellow lines in Figure 2b.

### Worsening Binding Affinity

When considering a past-to-present trend in viral collection date, there are significant correlations that show a worsening in antibody binding affinity in isolates collected from humans. See Figure 3. This worsening in binding affinity is not specific to particular antibodies. Though the antibodies’ performances are from independent distributions, as shown by the Kruskal-Wallis test in Supplementary Figure 4, the overall trend indicates that more recent isolates have mutated to better evade existing antibodies (antibodies that were previously elicited by vaccination or infection or developed for therapeutic usage).

**Fig. 3.**
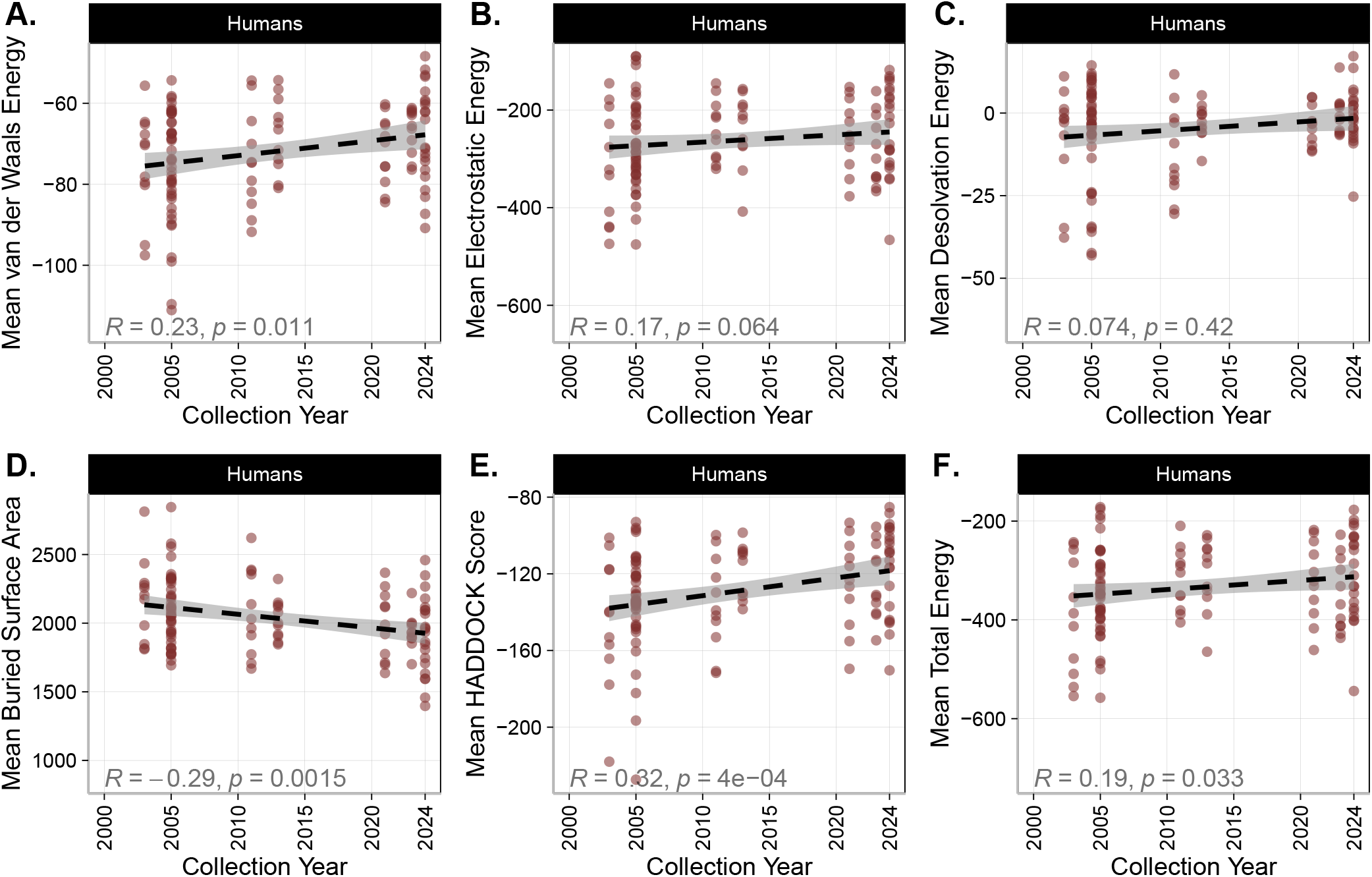
Antibody binding performance metrics over time for isolates collected from humans. Statistics shown are Spearman correlations. Overall, these plots show a worsening trend in most antibody binding metrics of the human samples.

As shown in Figure 4A, there is no overall trend in the graph edit distance of the isolates over time. In other words, when comparing to the 1959 isolate, EPI242227, interactions are not necessarily more or less abundant in more recent isolates than older ones overall.

**Fig. 4.**
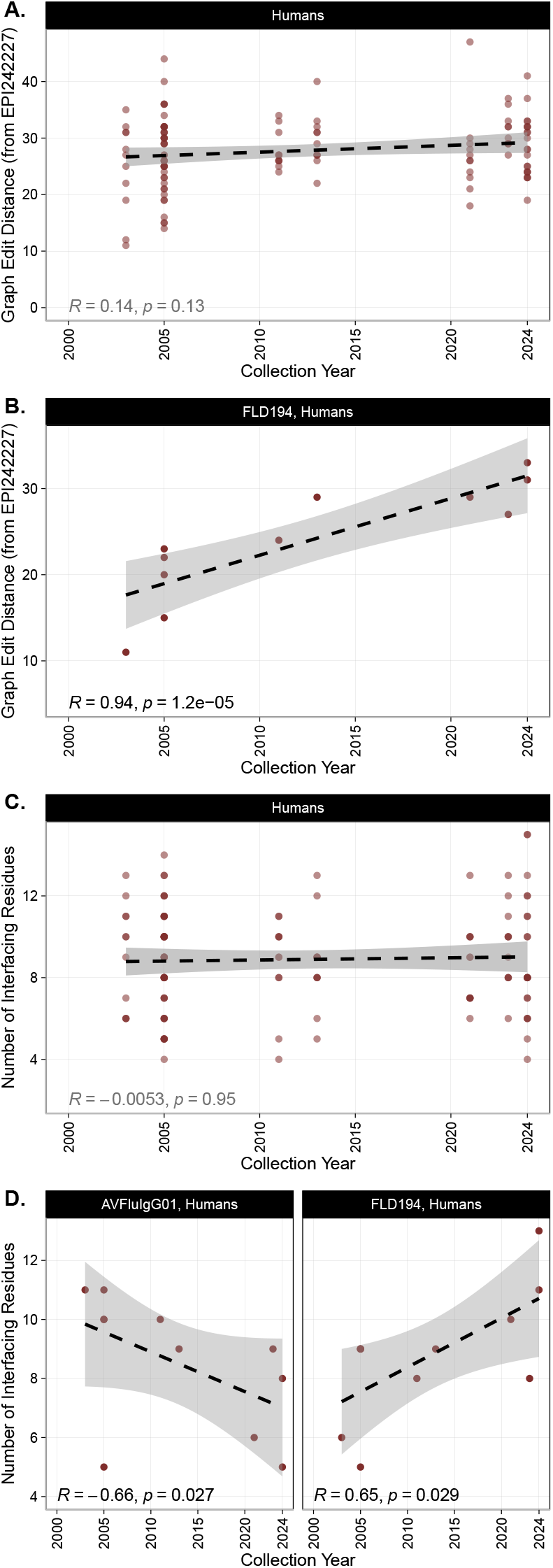
Graph-based analysis results showing the correlation of the graph edit distance and interfacing residue counts against collection year. Subfigures A and C show the results of all antibodies. Overall, graph edit distance did not change significantly over time. Subfigures B and D show Spearman correlations for specific antibodies of interest. Graph edit distance increased significantly in humans for FLD194. Number of interfacing residues increased in FLD194 in humans.

As shown in Figure 4C, we see a statistically significant decreases of interfacing residues over time in antibodies AVFluIgF01 against human isolates. Conversely, we see a statistically significant increase in interfacing residues in FLD194 against human residues. These results indicate an overall worsening in antibody affinity to more recent H5N1 isolates, which poses a risk to public health in that the virus may evade existing antibodies and risk the development of severe sickness in humans.

#### Mutational Effects

As indicated in Shi et al. (2014), there are various sites in the HA1 receptor binding domain that enable infection of mammals when mutated. Our results show several statistically significant differences in the binding affinity of antibodies given polymorphism at sites that allow mammalian infection.

Of note, N158S, T160A/S/V, E190N, and G225R all result in weakened antibody binding affinity across multiple metrics. Conversely, T160K and G228S increase binding affinity in some metrics. Significant changes based in Van der Waals energies and HAD-DOCK Scores are shown in Figure 6 and all other metrics are shown in Supplementary Figure 5.

In Figure 5, subfigures A and B represent the worst and best binding structures across the experimental results, isolates EPI168674 and EPI2429052, respectively. Though the epitope is different between isolates, note the variation in quantity of polar contacts within the respective complexes. Figure 5D is an example of modest binding affinity between an isolate EPI658567 and antibody 12H5. However, this improved binding affinity compared to other isolates is not due to the G225R mutation as this residue is not in the epitope of the antigen.

**Fig. 5.**
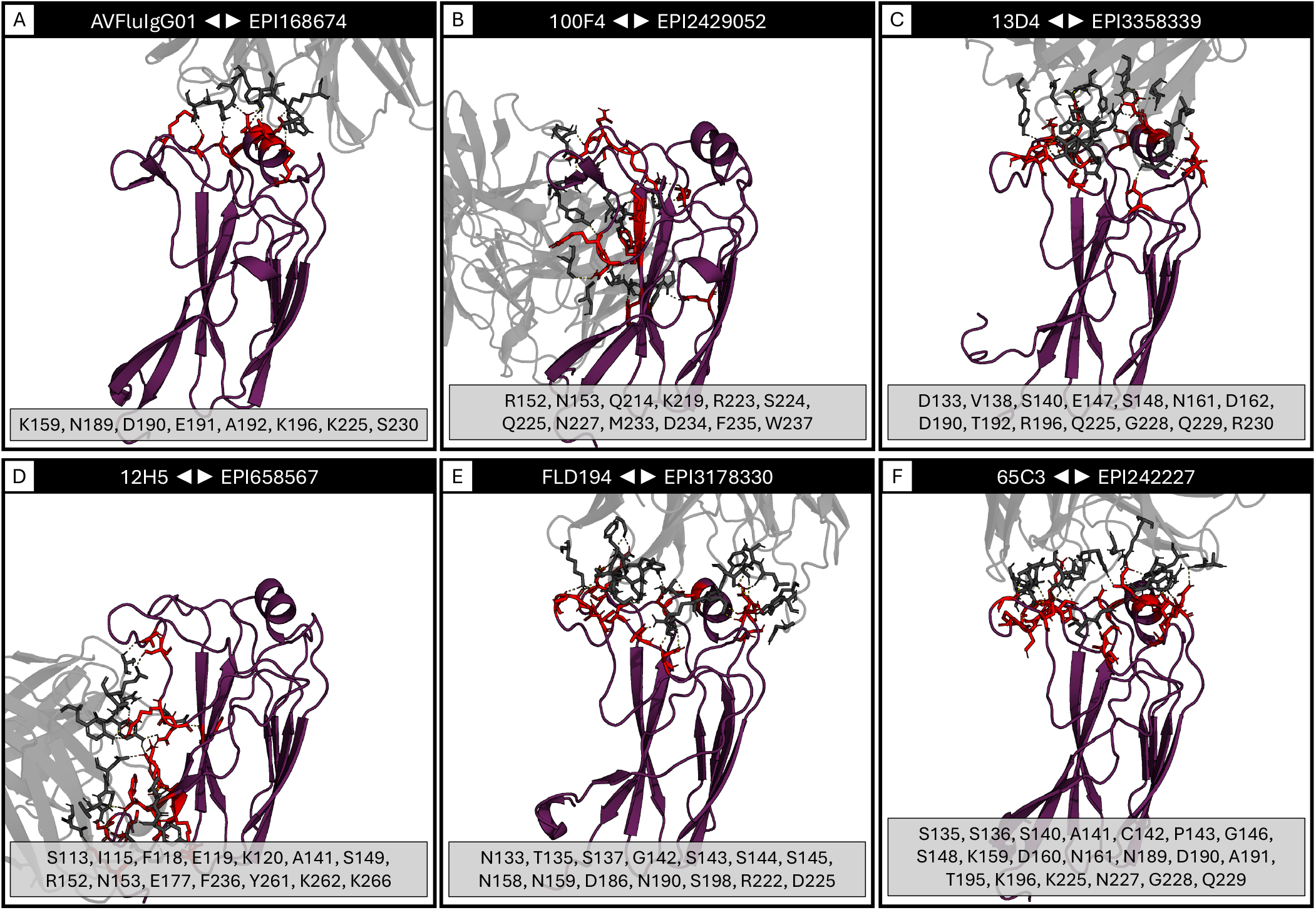
Example interface renderings showing the diversity in epitopes, residues, and binding affinity. The grey structure is the Fab portion of the docked antibody and the purple structure is the HA1 antigen with sticks designating the polar contacts between them. The list below each subfigure contains the interfacing residues on the antigen chain.

**Fig. 6.**
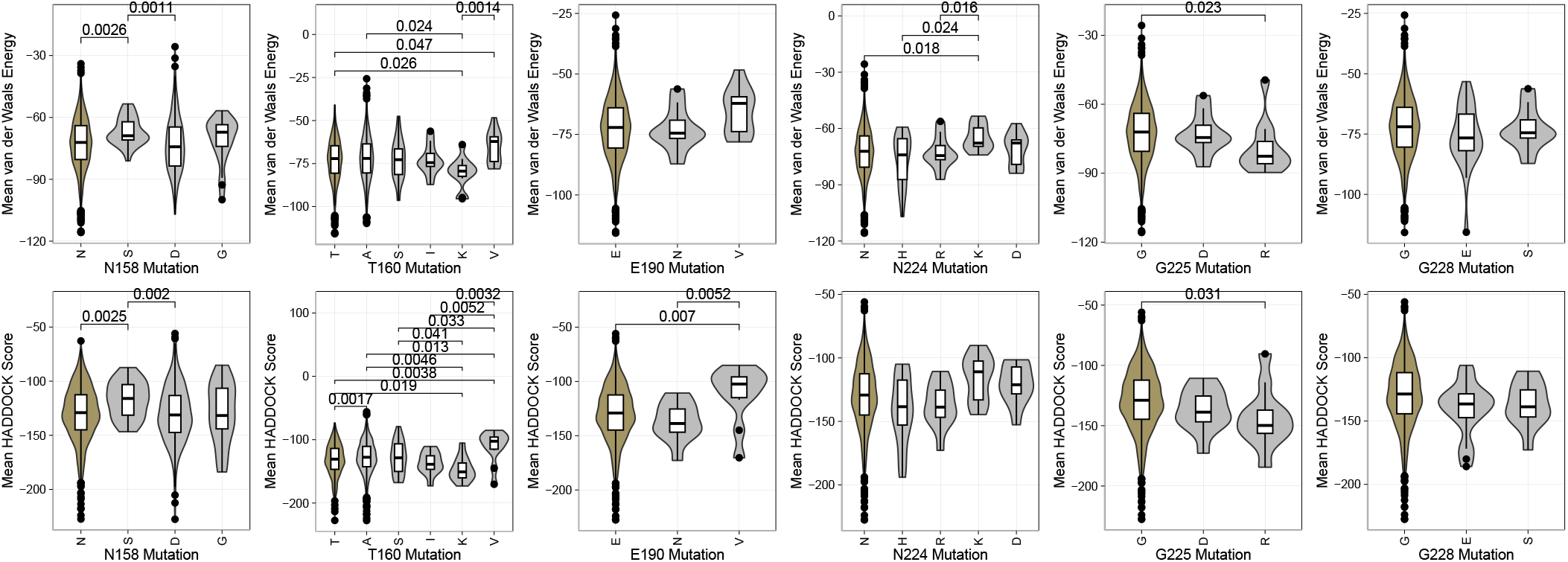
The distribution of Van der Waals energies and HADDOCK score docking metrics broken out by antigen mutations. The first amino acid shown on the left of each plot in gold represents the reference residue at that position as described in Shi et al. (2014). Statistical comparisons shown are significant Wilcoxon Rank Sum test p-values at the *α*<0.05 level.

Figure 5E shows the poor binding affinity of isolate EPI3178330 with antibody FLD194 due to the E190N mutation. However, this residue forms a polar contact with the antibody CDR loop. The mutation from glutamic acid (E), a negatively charged side chain, to asparagine (N), a polar uncharged side chain.

Isolate EPI242227 is the oldest isolate in the dataset, collected in 1959. Note that this isolate contains the N158D mutation and its interaction with various antibodies results in a wide range of Van der Waals energies from -52.81 (worst) to -92.84 (best, with antibody 65C3 as shown in Figure 5F).

#### Interfacing Residue Prevalence

An analysis of the interfacing residues in the best complexes from all 1,804 experiments shows patterns in particular residues forming polar contacts with the antibodies tested. Residues 156, 193, 222 are often interfaced (*≥*25% prevalence) in the antigen epitope. These are shown in red in Figure 7.

**Fig. 7.**
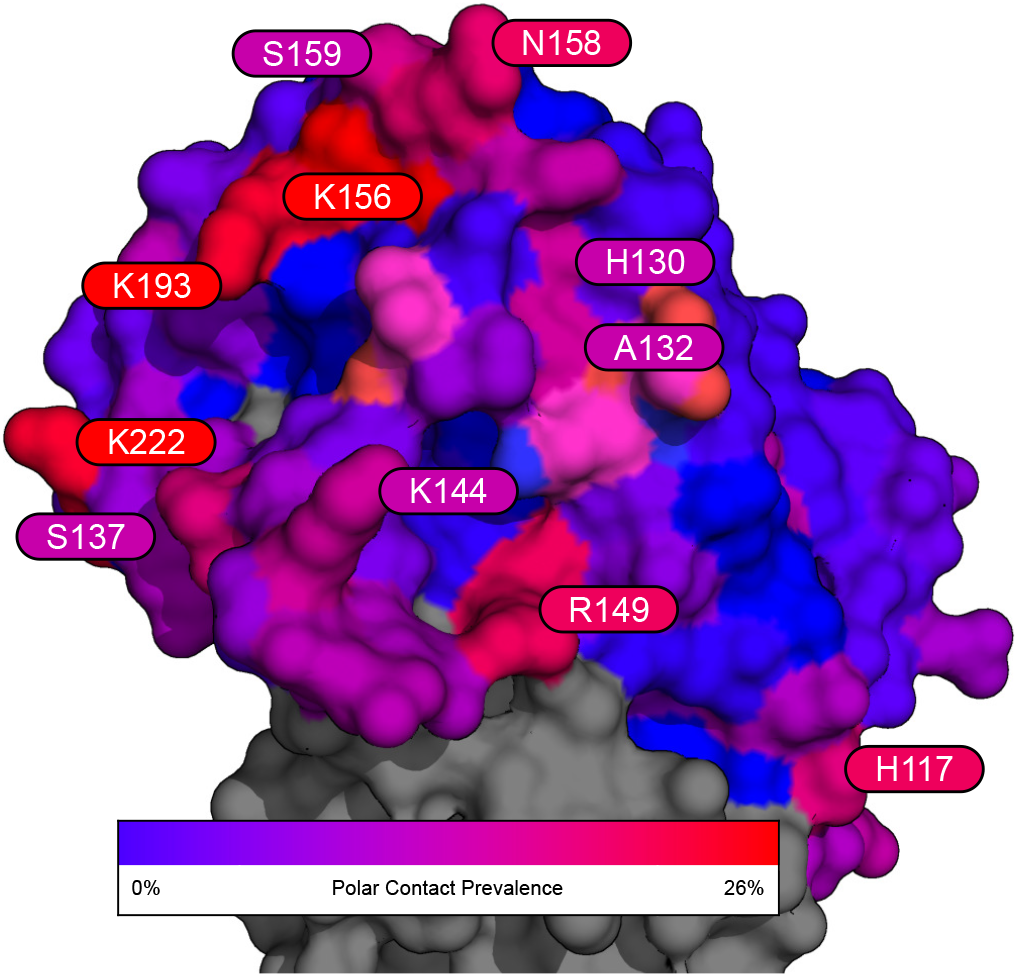
Surface rendering of the HA1 globular head domain (reference PBD 2FK0) showing the prevalence of each residue to form polar contacts (within 3Å of antibody residues) across the experiments in this study. Annotated residues are those with *≥*16% prevalence.

Many of the frequently interfaced residues shown in this study closely neighbor antigenic and receptor binding sites reported in Sriwilaijaroen and Suzuki (2012) Figure 2. Residues 130, 132, 158, and 159 are part of two glycosylation sites that, in 2012, were only found in seasonal H1 viruses (35).

## Discussion

### Relation to Prior Studies

This *in silico* analysis yields the trend of a reduction in binding affinity for neutralizing antibodies against H5N1-designated influenza isolates. This reduction in binding affinity reinforces previous studies that evolution has occurred to yield HA proteins that are elusive of antibody neutralization (62, 63). The trend observed is consistent with empirical studies and strengthens the novel *in silico* approach taken within this study. Current bio-surveillance efforts focus on critical mutations that have been shown to increase virulence or transmission risk, such as those included in the influenza risk assessment framework (IRAF) (64). This work suggests that the current arsenal of broadly neutralizing antibodies against H5N1 is becoming increasingly insufficient as H5N1 evolves, indicating a need for more studies to identify effective antibodies.

### Diversity of H5N1 Interactions

As shown in Supplementary Figure 3, high disparities in antibody binding affinity exist between different sequences designated as H5N1. For example, in Supplementary Figure 3, sequence EPI597824 has a large disparity in Van der Waals energy between antibody 3C11 and AvFluIgG03. In addition, Supplementary Figure 5 demonstrates the effect of a single aminoacid mutation on docking metrics. One change in an amino acid residue can lead to statistically significant differences in antibody binding affinity. These disparities highlight the need to continue to elucidate how differences in amino acid sequences alter binding affinity to various antibodies. Categorizing H5N1 influenza outside of the primary amino acid sequence but on functional binding analyses may yield more effective treatments in the future.

### Zoonosis Analysis

In Figure 2a, a notable transition pattern is observed from avian species (class Aves) to mammals (class Mammalia), likely attributable to a mutagenic drift. Recent empirical studies have investigated the mutational dynamics of H5N1, revealing changes in the hemagglutinin (HA) protein. While H5N1 influenza prefers binding to the *α*2-3 sialic receptors in birds, this study demonstrated a binding affinity of H5N1 to *α*2-6 sialic acid receptors, predominant in mammals, at almost equal proportion (4). These authors also show that mutations that decrease neutralization by sera from mice and ferrets immunized with the vaccine candidate reference strain A/American Wigeon/South Carolina/USDA-000345-001/2021 exist in some of the most recently collected mammalian samples, the dairy cow outbreak starting in April 2024 (4). Concurrently, we show here that mutations accumulated over time in the HA protein will confer reduced neutralization by antibodies more broadly than the current clade 2.3.4.4b H5N1 outbreak.

This *in silico* study aligns with these findings, indicating a progressive decrease in H5N1’s binding affinity to antibodies in our isolates over time. As illustrated in Figure 3, there is a marked decline in affinity for human isolates. As the virus diversifies in the avian populations, the potential pool of strains with zoonotic potential to infect mammals increases. The phylogenetic and transmission analysis show much more frequent transmission from avian populations to mammalian populations. This result indicates that much of the evolution is occurring in birds. This suggests an evolutionary trajectory in birds for H5N1 towards increased infection in mammalian hosts with a concomitant immune evasion of the virus in mammals.

#### Isolate EPI3358339

EPI3358339, a H5N2 subtype isolate, was added to this study as it is from the recent human infection of H5N2 avian influenza seen in Mexico. Unfortunately, this strain was found in a person from Mexico who died of complications due to the infection. However, it is not yet known if the zoonotic “jump” the isolate is cause for concern or if the individual had other comorbidites that played a role in his death. It is also not known if this case is related to recent poultry outbreaks in the area.

The experimental docking conformation between antibodies and this antigen (antibody 65C3 shown in Figure 5F) are predicted to have a relatively strong binding affinity (e.g., Van der Waals energy: [-51.96, -95.45]) across all the tested antibodies.

Thus, our experiments using this isolate’s HA do not indicate that this isolate is highly mutated, though some mutations may have reduced the Van der Waals and electrostatic energies of the interaction with this individual’s existing antibodies, if any.

### Structural Analysis

When comparing the predicted docking outputs to empirical structures, such as those listed in Table 1, we see similar binding epitopes in the the predicted docking complexes versus the empirical structures (42–49).

Furthermore, the binding conformations seen in the empirical structures often mimic the predicted complexes in the experiments in this study, though various mutations affect the binding angle, polar contacts, electrostatics, and overall affinity. These results support the confidence in the predictive accuracy of the *in silico* experiments given their similarity to empirically derived structures. However, empirical studies are still needed to validate specific complexes.

### Motifs of Interest and Future Research

H5N1 exists as an endemic in avian populations. This endemic creates a pool containing vast numbers of host species in which the RNA virus evolves rapidly. It is an endemic that is difficult to diminish due to the nature of the HA protein’s host receptors lacking homogeneity. Serological immunity from vaccination or prior infection in avian hosts may have yielded selective pressures in the evolution of specific mechanisms of entry for H5N1 (65). Subsequently, high infection rates lead to new mechanisms of entry. Over time, serological immunity from original vaccination and/or infection is reduced, and the cycle of influenza transmission continues.

Highly conserved portions of HA are of high interest. The three primary conserved elements of the receptor-binding site (RBS) on the HA1 subunit are the 130 and 220-loops and the 190-helix (66–68). As shown in Figures 5 and 7, the antibodies docked to conserved motifs on the studied H5N1 strains, further supporting the empirical literature that initially identified their neutralizing capability.

More recent development of multivalent mRNA-based vaccines has been successful in H5 influenza A clade 2.3.4.4b (from which there are 15 sequences used in the structural aspects of this study) (69). The selection of high-quality sequences that elicit strong antibody responses is a complex process in mRNA vaccine development. However, *in silico* modeling, as presented in this study, reduces the wet laboratory workload to evaluate candidate sequences from which vaccines can be developed.

In addition, our broad analyses of various antibodies versus strains of interest may guide future therapeutic antibody development. Antibodies tested within this study, particularly those with high affinity to studied strains that may bear high homology to those that will arise in the future, can be used as a basis for future pharmaceutical development.

Computational modeling of immunoprotein interactions as shown in this study and previous works (24–29) have proven to be highly effective in the prompt prediction and understanding of the health impacts of pathogen variants. For H5 influenza, this study, along with recent preprints (18, 70), show an overall trend of worsening antibody binding and depicts the recent increase in avian-to-mammalian transmissions due to various mutations. This suggests that there is an impending danger to human health for highly pathogenic strains of H5 influenza that can infect avian and mammalian livestock and jump to humans.

More broadly, these results indicate that the virus has potential to move from epidemic to pandemic status in the near future. “Pandemic” here refers to the geographic spread of a virus, which H5N1 has already achieved, but these results more specifically assert that the worsening trend of the antibody performance along with the already present animal pandemic is a cause for concern for an eventual human pandemic.

## Contributors

Authors SY, KO, and SGM retrieved the H5 antigen sequences from GISAID and antibody structures from Protein Data Bank. SY performed the structure prediction of the antigens. Author RAW performed the clustering analysis. Authors KO and CTF curated the metadata of the H5 sequences. Author DJ performed the phylogenetic analyses. Authors SY and CTF performed the protein structure analyses. Author PJT performed the multiple sequence alignment. Author RJ3 performed the graph-edit distance analysis. Author CTF performed the docking experiments and statistical analyses and generated all figures. All authors wrote and reviewed the manuscript.

## Acknowledgements

We gratefully acknowledge all GISAID data contributors (i.e., the authors and their originating laboratories) responsible for obtaining the specimens, and their submitting laboratories for generating the genetic sequence and metadata and sharing via the GISAID Initiative, on which this research is based.

We acknowledge the following entities at the University of North Carolina at Charlotte: Academic Affairs, The Office of Research, The Center for Computational Intelligence to Predict Health and Environmental Risks (CIPHER), The Department of Bioinformatics and Genomics, The College of Computing and Informatics, and the University Research Computing group. We gratefully acknowledge the support of the Belk Family.

## Data Sharing Statement

All code, data, results, and additional analyses are openly available on GitHub at: https://github.com/colbyford/Influenza_H5-Antibody_Predictions. These data include all sequences and folded structures for the isolates and antibodies used in this study, analysis scripts, and docking metrics.

## Supplementary Information

**Fig. 1.**
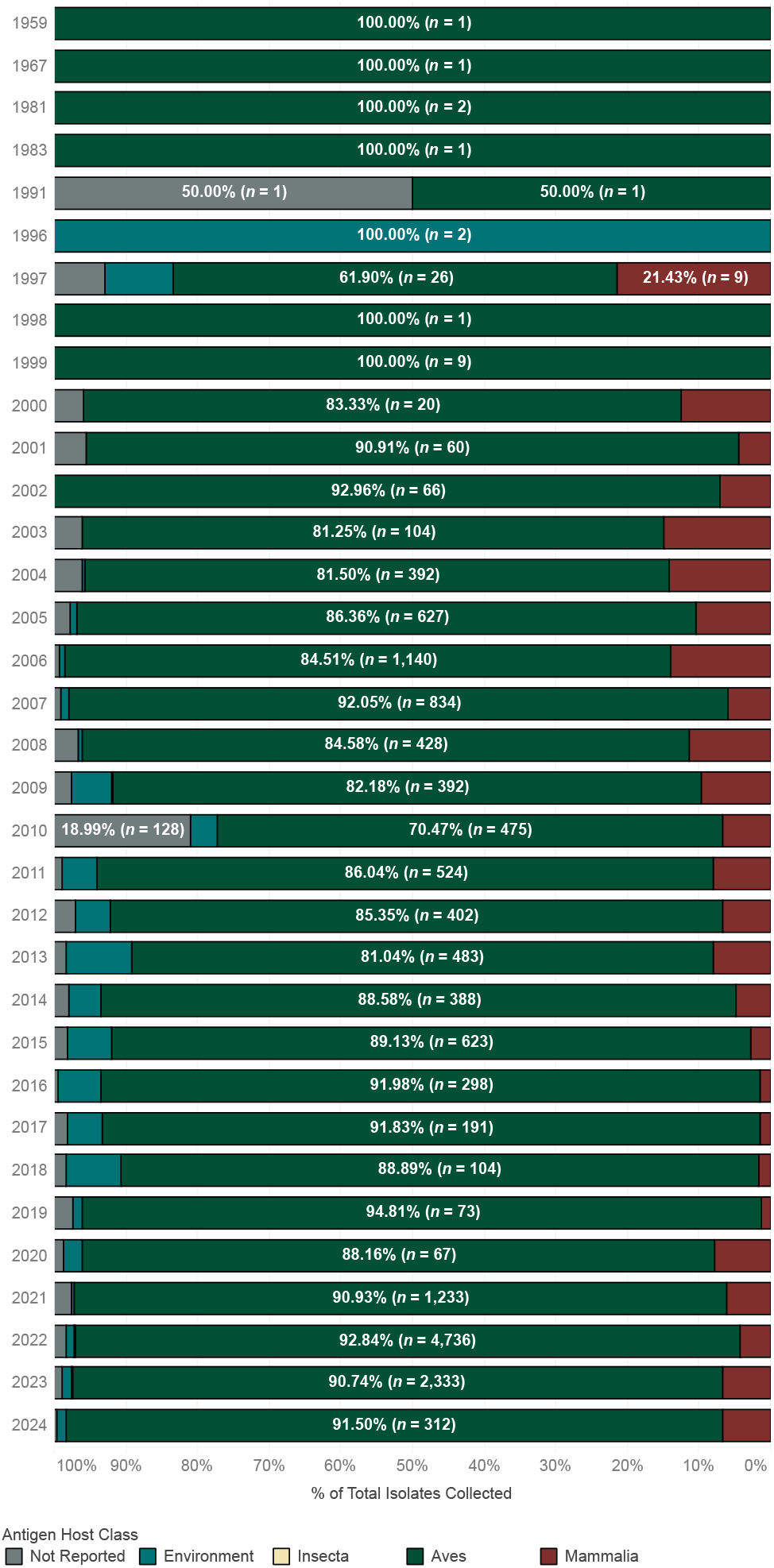
Bar chart of the proportion of isolates collected from hosts of various taxonomic classes by year.

**Fig. 2.**
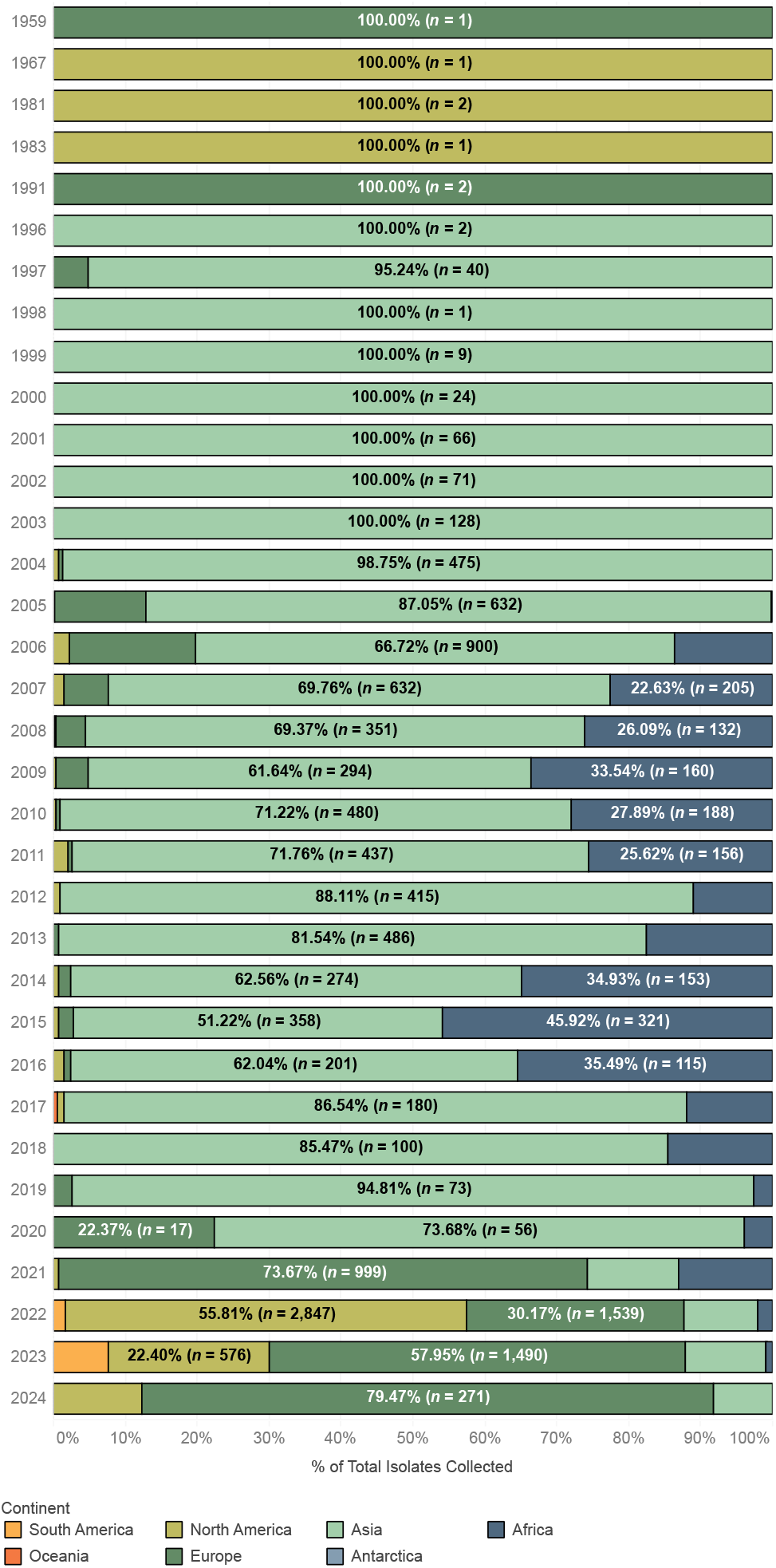
Bar chart of the proportion of isolates collected from each continent by year.

**Fig. 3.**
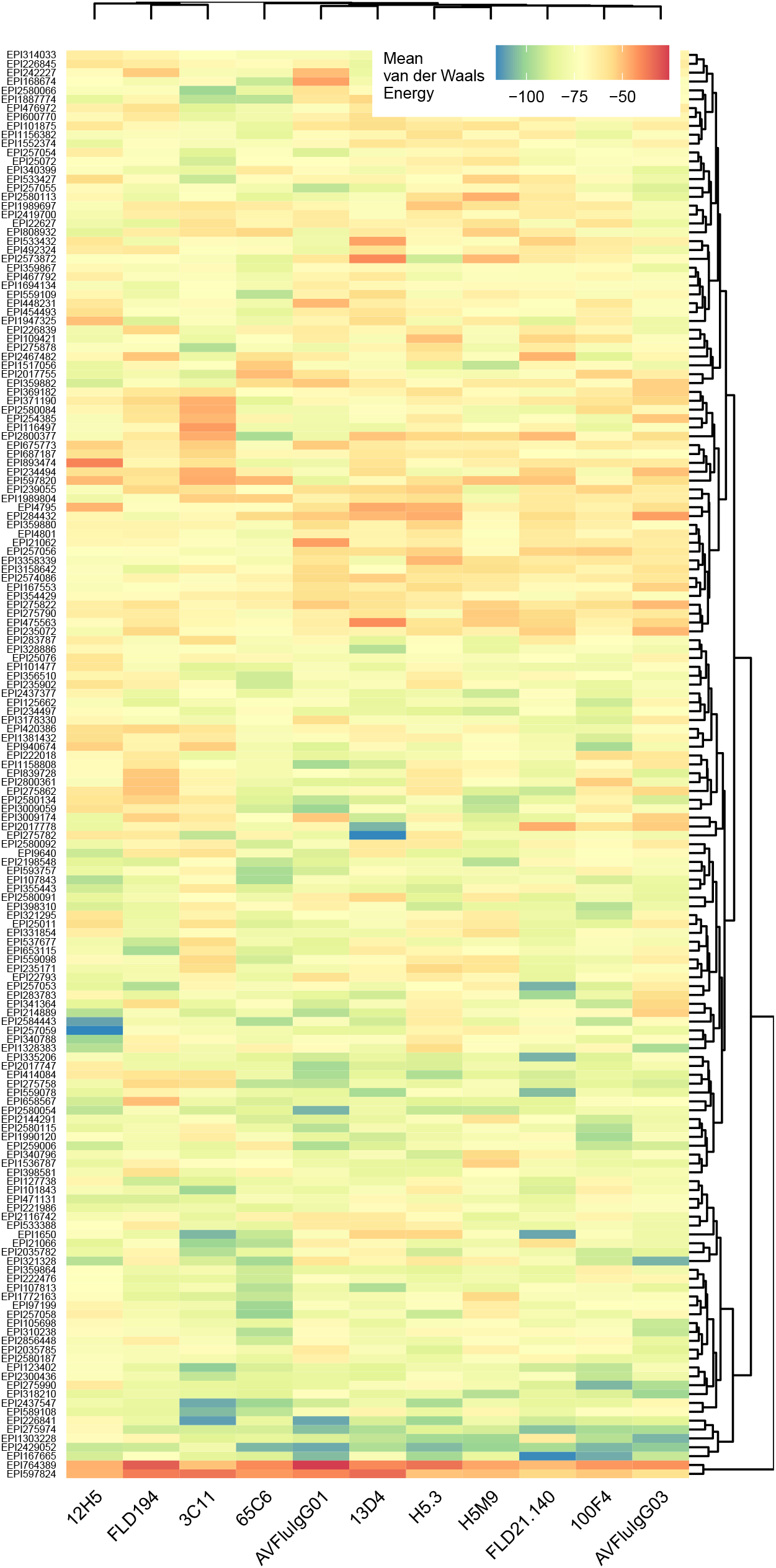
Heatmap of van der Waals energies for each antibody-antigen experiment, shown with cluster dendrograms.

**Fig. 4.**
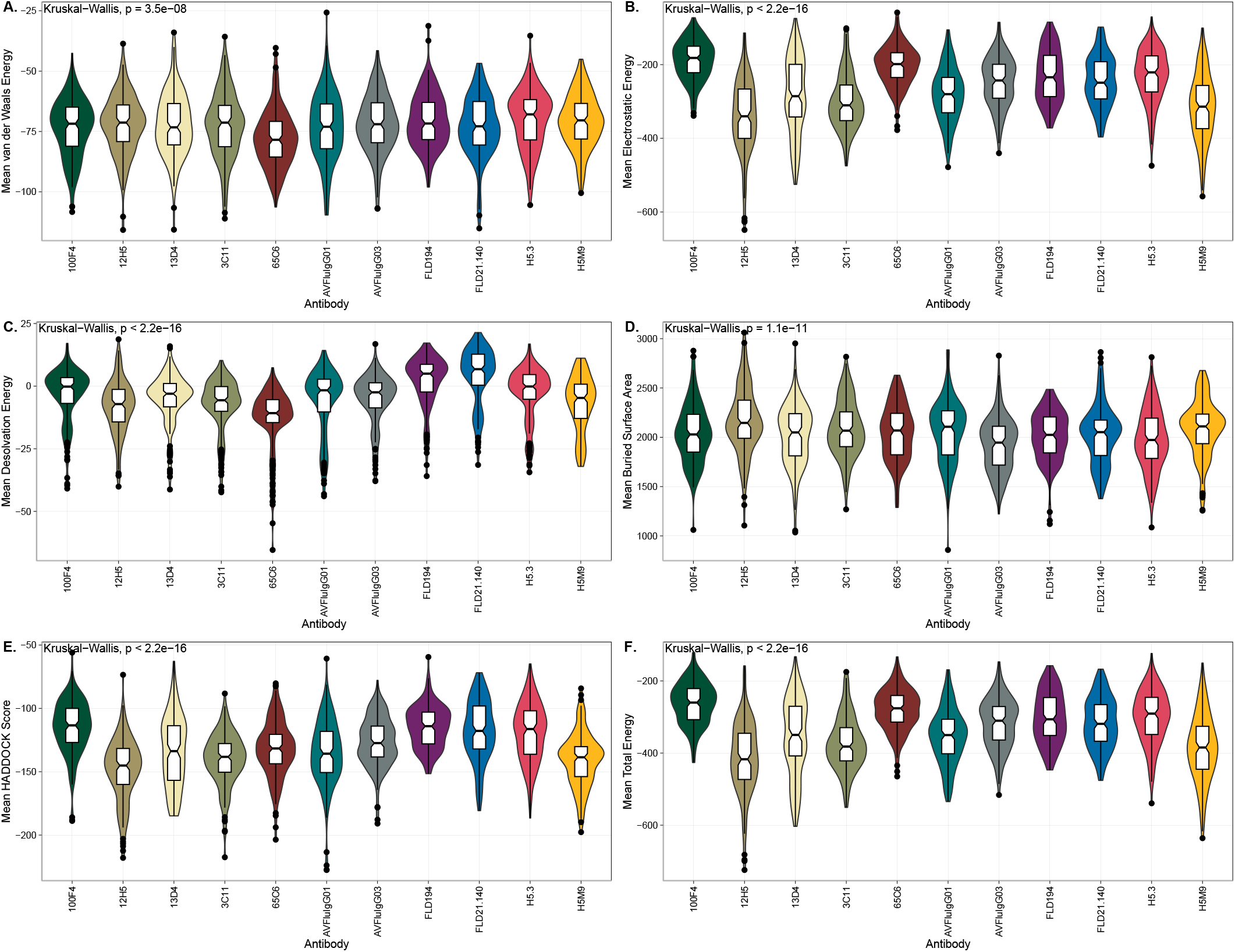
The distribution of various docking metrics broken out by antibody. Statistics shown are Kruskal-Wallis test p-values.

**Fig. 5.**
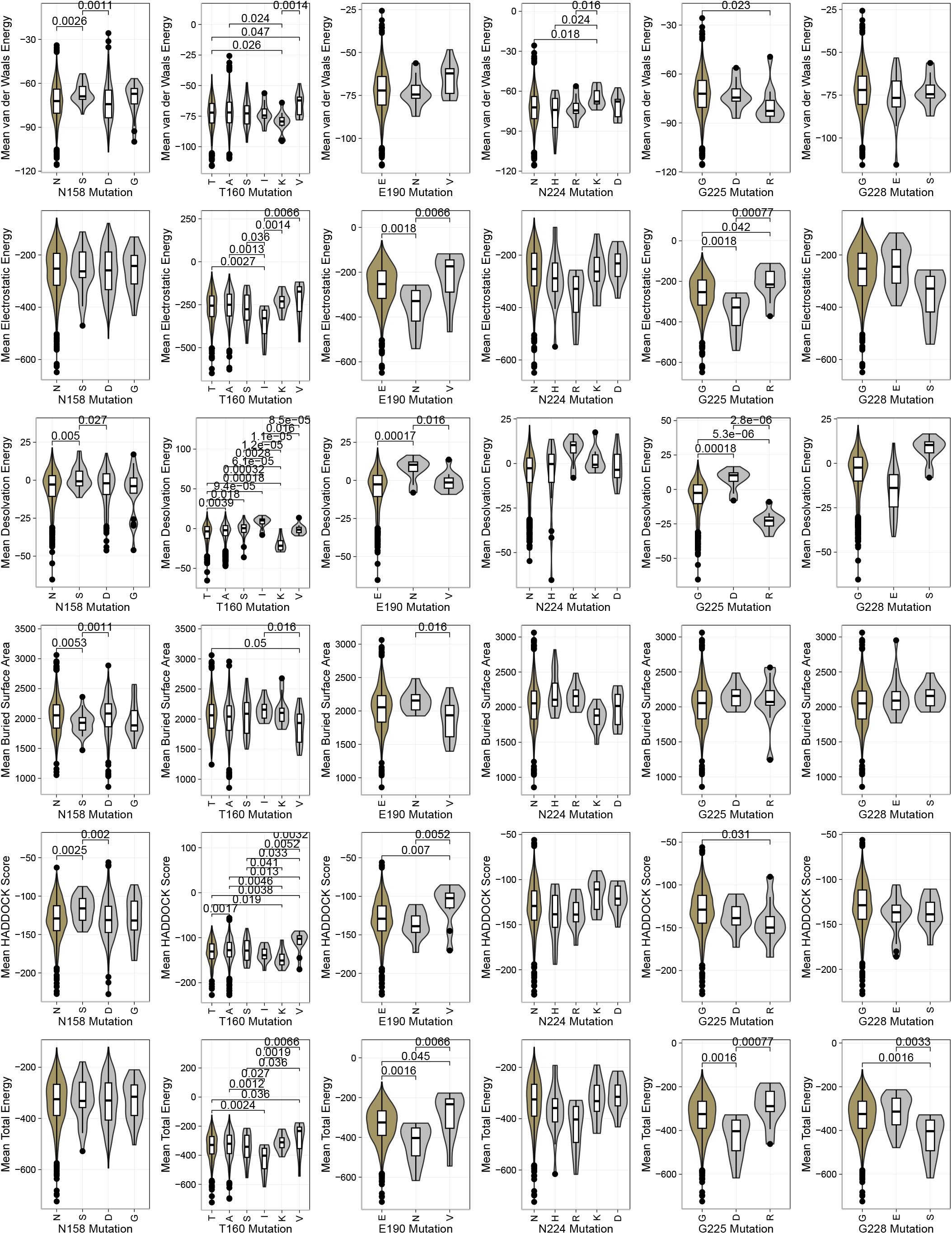
The distribution of various docking metrics broken out by antigen mutations. The first amino acid shown on the left of each plot in gold represents the reference residue at that position as described in Shi et al. (2014). Statistical comparisons shown are significant Wilcoxon Rank Sum test p-values at the *α*<0.05 level.

## Footnotes

1 Predicted local distance difference test (pLDDT): An estimate of local confidence, scaled from [0, 100], where higher scores indicate higher confidence in the protein conformation.

2 Container GitHub Repository: https://github.com/colbyford/HADDOCKer

3 Docker Hub Images: https://hub.docker.com/r/cford38/haddock

